# Nurturing diversity and inclusion in AI in Biomedicine through a virtual summer program for high school students

**DOI:** 10.1101/2021.03.06.434213

**Authors:** Tomiko Oskotsky, Ruchika Bajaj, Jillian Burchard, Taylor Cavazos, Ina Chen, Will Connell, Stephanie Eaneff, Tianna Grant, Ishan Kanungo, Karla Lindquist, Douglas Myers-Turnbull, Zun Zar Chi Naing, Alice Tang, Bianca Vora, Jon Wang, Isha Karim, Claire Swadling, Janice Yang, AI4ALL Student Cohort 2020, Marina Sirota

**Affiliations:** Department of Pediatrics, UCSF, San Francisco, CA, USA; Bakar Computational Health Sciences Institute, UCSF, San Francisco, CA, USA; Department of Bioengineering and Therapeutic Sciences, UCSF, San Francisco, CA, USA; Cognitive Science and Computer Science Programs, UCLA, Los Angeles, CA, USA; Program in Biological and Medical Informatics, UCSF, San Francisco, CA, USA; Pharmaceutical Sciences and Pharmacogenomics, UCSF, San Francisco, CA, USA; Berkeley Institute for Data Science, UC Berkeley, Berkeley, CA, USA; School of Medicine, UCSF, San Francisco, CA, USA; Department of Epidemiology & Biostatistics, UCSF, San Francisco, California, USA; Institute for Neurodegenerative Diseases, UCSF, San Francisco, CA, USA; Quantitative Biosciences Consortium, UCSF, San Francisco, CA, USA; QBI COVID-19 Research Group (QCRG), San Francisco, CA, USA; Quantitative Biosciences Institute (QBI), UCSF, San Francisco, CA, USA; J. David Gladstone Institutes, San Francisco, CA, USA; Department of Cellular and Molecular Pharmacology, UCSF, San Francisco, CA, USA; Department of Bioengineering, UC Berkeley, Berkeley, CA, USA; Saint Francis High School, Mountain View, CA, USA; AI4ALL, Oakland, CA, USA; Canton High School, Canton, MI, USA; Department of Computer Science, Stanford University, Stanford, CA, USA

## Abstract

Artificial Intelligence (AI) has the power to improve our lives through a wide variety of applications, many of which fall into the healthcare space; however, a lack of diversity is contributing to flawed systems that perpetuate gender and racial biases, and limit how broadly AI can help people. The UCSF AI4ALL program was established in 2019 to address this issue by promoting diversity and inclusion in AI. The program targets high school students from underrepresented backgrounds in AI and gives them a chance to learn about AI with a focus on biomedicine. In 2020, the UCSF AI4ALL three-week program was held entirely online due to the COVID-19 pandemic. Thus students participated virtually to gain experience with AI, interact with diverse role models in AI, and learn about advancing health through AI. Specifically, they attended lectures in coding and AI, received an in-depth research experience through hands-on projects exploring COVID-19, and engaged in mentoring and personal development sessions with faculty, researchers, industry professionals, and undergraduate and graduate students, many of whom were women and from underrepresented racial and ethnic backgrounds. At the conclusion of the program, the students presented the results of their research projects at our final symposium. Comparison of pre- and post-program survey responses from students demonstrated that after the program, significantly more students were familiar with how to work with data and to evaluate and apply machine learning algorithms. There was also a nominally significant increase in the students’ knowing people in AI from historically underrepresented groups, feeling confident in discussing AI, and being aware of careers in AI. We found that we were able to engage young students in AI via our online training program and nurture greater inclusion in AI.

## Introduction

Artificial Intelligence (AI) has the power to improve our lives through a wide variety of applications. Just a few examples of how AI is being used to enrich our lives include search engines, autonomous vehicles, and facial-recognition, route-planning, and ride-hailing programs (1). The applications of AI to the biomedical, translational, and clinical realms are diverse ranging from discovering biomarkers and repurposing therapeutics, to improving disease diagnosis and automating surgery (2). Moreover, AI can help realize the promise of personalized medicine, a healthcare approach that aims to tailor medical decisions and treatments to individuals based on their intrinsic (e.g., genomic, age, sex) and extrinsic (e.g., diet, environmental exposures) factors (3).

Yet a lack of diversity can adversely affect how broadly AI will help people (4). For instance, if machine learning (ML) algorithms to diagnose skin cancer lesions were trained on data that largely represent fair-skinned populations, then the algorithms, no matter how advanced, would not perform as well on images of lesions in skin of darker color (5). We need diversity not just in the data we use in AI but also in the people working and leading in the field of AI. Currently, AI professors are mostly male (>80%), and among AI researchers, only 15% at Facebook and 10% at Google are female (6). Moreover, black workers represent 4% of the workforce at Facebook and Microsoft, and only 2.5% of Google’s entire workforce (6).

The UCSF AI4ALL program, established in 2019 and co-directed by Marina Sirota, PhD and Tomiko Oskotsky, MD, strives to promote greater diversity and inclusion in the field of AI in biomedicine, and to inspire tomorrow’s leaders to think about and know AI and to use AI ethically. UCSF AI4ALL recruits high school students from backgrounds underrepresented in AI, including females and students from minority racial and ethnic backgrounds, as well as students from low income families and those who are the first in their families to go to college. Through this tuition-free three-week summer training program, students gain experience with AI with a focus on applications to biomedicine, interact and work with a diverse set of role models in AI, including women and people of Black or African American background and Hispanic or Latino background, and learn about how AI can advance health. They receive broad exposure to AI topics through faculty lectures, and gain in-depth research experience through hands-on projects. Mentoring and career/personal development sessions with faculty, researchers, industry professionals and undergraduate and graduate students further enable personal growth and an opportunity to explore career interests at the intersection of Computer Science and Biomedicine. Due to the COVID-19 pandemic, the UCSF AI4ALL program held in 2020 shifted from an in-person, commuter program to a synchronized, online one with all the student research projects focusing on leveraging AI to advance our knowledge and understanding of COVID-19. Here, we provide an overview of the 2020 UCSF AI4ALL virtual summer program, share details about the research projects our students engaged in, and discuss the results of our program.

## Methods

We reviewed all 89 complete applications that were submitted to our program during the application period in March 2020, and assessed each candidate holistically prior to offering acceptances into our program.

The program itself was a three week program, which was held virtually in 2020. The first week focused on teaching the high school participants basic programming in python and introductory topics in machine learning and the second and third weeks focused on research projects, which in 2020 were all leveraging AI to COVID-19. Google CoLab notebooks were used as a simple means to share code with students and run Python. Each morning began with time for students to ask questions as well as participate in our ice-breaker activities. The program had daily guest lectures by diverse faculty from UCSF focused on application of AI in biomedicine and covering a wide range of topics from clinical data analysis, to diagnostic and therapeutic strategies leveraging molecular measurements. There were also panels composed of UCSF AI4ALL student alumni, undergraduate students, graduate students, and professionals from private companies with backgrounds in AI within biomedicine and other disciplines. Our panelists, many of whom were women and people from diverse racial and ethnic backgrounds, spoke with our students and shared insights about their work and their journeys. Each week, our Alumni TAs led Community Building Session engaging the class of students in fun, bonding exercises. We also held a personal growth session to develop the students’ communications skills. The end of the program symposium included student presentations on their research projects as well as a Keynote talk on AI in Biomedicine. A copy of the 2020 program schedule is available https://ai4all.ucsf.edu/assets/2020_UCSF_AI4ALL_Program_Schedule.pdf

Students were asked to complete a survey at the beginning (Pre-) and at the conclusion (Post-) of our program. Mann Whitney U (MWU) test with continuity correction was used to compare Pre-to Post- survey responses (since surveys were anonymous, we could not compare these using tests designed for paired data), and to compare 2019 to 2020 Pre- and Post- survey responses. Bonferroni corrections were employed, and a significance threshold of 0.05 was applied to the results.

## Results

Of the 89 high school students who submitted applications to our program and the 38 applicants we accepted into the program, 29 enrolled in and completed the program.

All 29 students were females who were rising sophomores (21%), juniors (45%) or seniors (34%) in high school. Most of the students were from California (79/%), although several were from other states. The racial backgrounds of the students included Asian inclusive of those from the Indian subcontinent and Philippines (79%), Native Hawaiian or Other Pacific Islander/Original Peoples (3%), and Hispanic or Latino (7%), and 14% declined to state. Twenty-one percent will be first generation college students. (**Table 1**).

**Table 1.** Demographic characteristics of accepted students in the 2020 UCSF AI4ALL Summer program

| Characteristic | # students (total accepted: 29) | % students |
| --- | --- | --- |
| **Gender** |  |  |
| She/Her | 29 | 100% |
| He/Him | 0 | 0% |
| They/Them | 0 | 0% |
| **Race \*more than one category may be checked** |  |  |
| Asian (including Indian subcontinent and Philippines) | 23 | 79% |
| Black or African American | 0 | 0% |
| Native Hawaiian or Other Pacific Islander (Original Peoples) | 1 | 3% |
| Hispanic or Latino (including Spain) | 2 | 7% |
| White (including Middle Eastern) | 0 | 0% |
| Decline To State | 4 | 14% |
| **Grade Level Next Year** |  |  |
| Senior / 12th grade student | 10 | 34% |
| Junior / 11th grade student | 13 | 45% |
| Sophomore / 10th grade student | 6 | 21% |
| Freshman / 9th grade student | 0 | 0% |
| **Qualify for Free Lunch at School** |  |  |
| Yes | 4 | 14% |
| No | 25 | 86% |
| **1st Gen College Student** |  |  |
| Yes | 6 | 21% |
| No | 23 | 79% |
| **Home State** |  |  |
| California | 23 | 79% |
| Other | 6 | 21% |

### 1st week: Lessons in Python and Machine Learning

In the first week of the program, students spent the afternoons learning about machine learning concepts and programming in Python. We had seven UCSF graduate student instructors and teaching assistants (TA) to help with teaching during the first week. iPython notebooks with the in-class exercises were shared the evening before the class, to give students an opportunity to practice on their own before the solutions were reviewed in class.

#### Python Workshops

Students covered the basics of programming, data management, and data visualization in the first two days to prepare to code in Python language and work with data within a Google CoLab environment in preparation of their projects. Topics covered include programming basics (data types, logic, loops, functions), data structures, common Python packages, plotting with matplotlib, and using sklearn. During the lesson, students were placed in breakout rooms with teaching assistants to review coding exercises and practice programming activities together.

#### Lessons on Machine Learning

Topics covered in ML include Data and Bias, Clustering, Classification, Naive Bayes, Regression, and Neural Networks. To facilitate remote instruction, we employed a reverse classroom paradigm, in which the instructors produced a 15-20 minute lecture video to be watched before each classroom session. The general structure of live sessions include 15 minutes of topic review first, then the rest of the session covering either conceptual activities or reviewing and practicing ML exercises on CoLab notebooks. For activities, students were placed into breakout rooms with a teaching assistant. Since students come from various backgrounds of ML and programming familiarity, collaboration within the breakout rooms were encouraged.

The instruction was carried out in an inverse classroom setting where the participants could ask the instructors and TAs questions after having watched the lectures.

### 2nd and 3rd weeks: Research Projects and Presentations

#### Applying AI to COVID-19

The students were assigned to one of five groups based on their preference, each working on a research project that applied AI to the characterization, classification, or prediction of COVID-19 leveraging different types of biomedical data - gene expression, proteomics, imaging and clinical. Each project team was led by a UCSF graduate student, medical student, and/or postdoctoral scholar and co-led by an alumni TA.

On the last day of the program, students shared findings from their group’s research project during our Final Symposium. Each presentation was approximately 15 to 20 minutes in length, with time for questions, and each student presented a portion of their group’s work. The event was attended virtually by over 100 people, including faculty, graduate students and postdoctoral scholars from UCSF and other institutions, program participants and their invited family members. A videorecording of the Final Symposium, including our Keynote Speaker’s talk and the students’ presentations is available, https://youtu.be/uImjiHl7MDw.

#### Project 1: AI for Global Health - AI and COVID-19 Time Series Diagnosis Data

Students learned how to develop machine learning algorithms with utility for lower middle income country (LMIC) settings. Their objective was to develop an algorithm that can predict the number of cases in a given country. Students used publicly available daily time series data describing confirmed COVID-19 infections and deaths per country and states across the world (over 266 regions) aggregated from the Johns Hopkins Center for System Sciences downloaded on July 1, 2020 (https://github.com/CSSEGISandData)(7). Each student then manually pre-processed the dataset to a format in which they could conduct exploratory data analysis. The educational approach was to allow students to have first-hand experience in discovering the optimal way to plot and analyze various features of the data they were working with by experimenting with different visualization libraries and troubleshooting together real-time through video conferencing. Recognizing the diversity in time series trends between countries during exploratory data analysis, students chose to narrow the scope of the problem to focus on a specific country, selecting India due to its large number of cases and disparity in public health services. Students then did a literature review to understand the public health issues in India and how to design an algorithm that may actually provide utility to key stakeholders in the region.

Visualizing the trend of confirmed infections in India, they decided to develop a forecasting algorithm that can aid in identifying how many resources a given country or state will need. Students were then presented with high-level information on several ML techniques used for time series data analysis, such as autoregression (8), Holt-Winters (9) exponential smoothing, and neural networks (10). Following a group debrief, students were allowed to select modeling approaches that interested them. Afterwards, they trained, developed, and tested three different algorithms: autoregression, feed-forward neural network, and Long Short-Term Memory (LSTM) recurrent neural network (**Fig 1a,b**).

**Fig 1.**
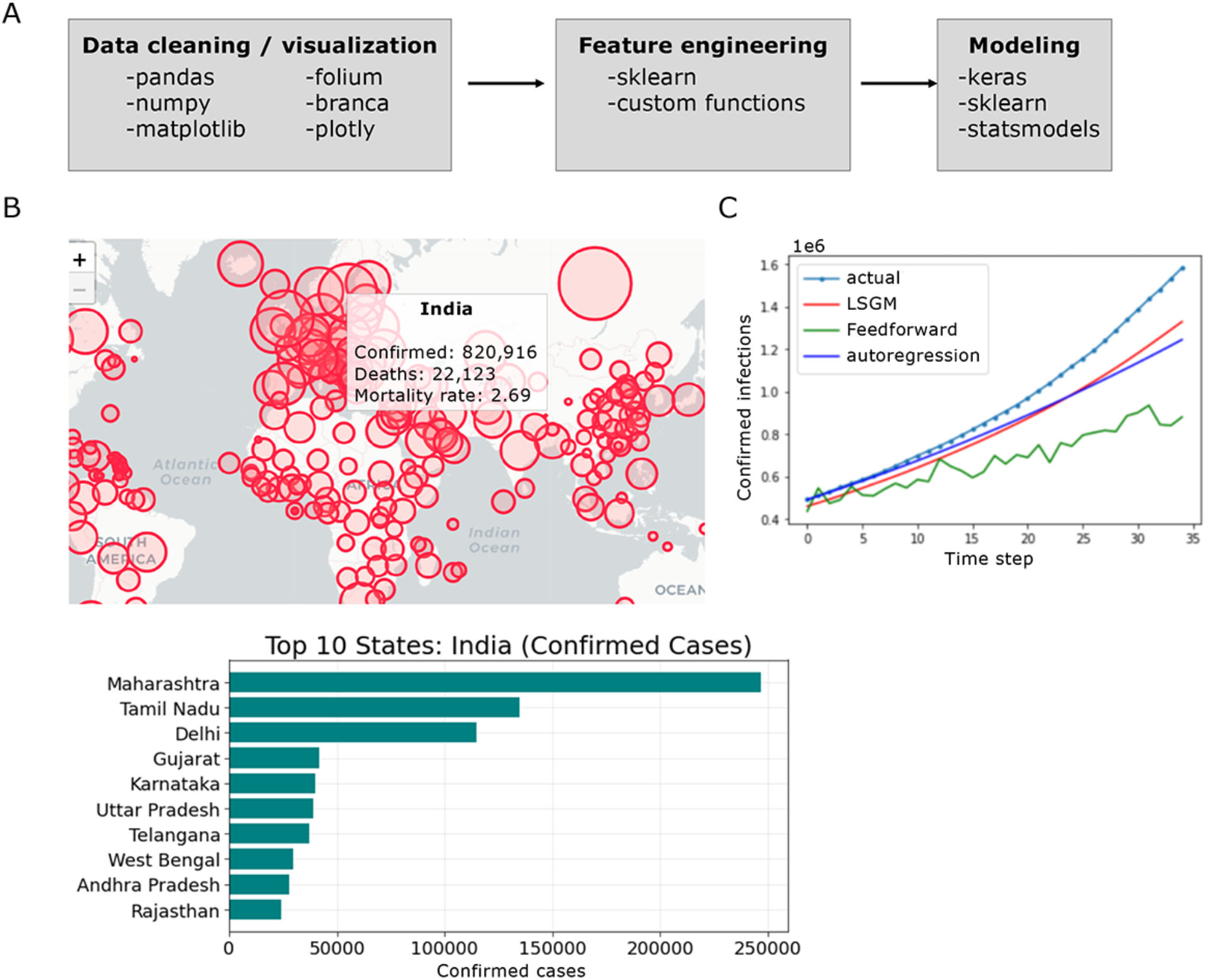
AI for Global Health. **A.** Schematic of major ML skills explored with packages/utilities used for instruction. **B.** Examples of data visualizations created by students. **C.** Model predictions compared to actual India COVID-19 data. Mean average percent error was 8.23%, 10.08%, 82.35% for autoregression, LSTM, and feedforward neural networks respectively.

Students first started with a simple ML technique for time series data, known as autoregression. Afterwards, they decided to see whether it was possible to leverage data from other countries that may be useful in the same prediction. They developed a feedforward neural network algorithm that leveraged data from 266 other countries/regions and predicted the most recent 5 days of COVID-19 cases in India. Students discovered that this model performed worse than the simple ML technique (Mean Average Percentage Error: 82.35% for feedforward neural network vs. 8.23% from autoregression) (**Fig 1c**). The students became interested in trying LSTM recurrent neural networks, due to their unique ability to model time-series data better than feed-forward networks. They trained the model to predict the next 5 days of data from the most recent 15 data points, and found it performed slightly better than the feed-forward, but not as well as the simple ML technique (Mean Average Percent Error: 10.08% for LSTM) (**Fig 1c**).

#### Project 2: AI and Proteomics - COVID-19 Protein-Protein Interactions (PPI) predictions

Students learned to implement supervised learning techniques to predict host protein interactors, given primary amino acid sequences of various viral proteins, and to test if proteins of similar sequences would interact with the same host proteins. The dataset was curated from two publicly available sources - 1) a host-pathogen protein-protein interaction (PPI) data in HEK293T cells for HIV (11), HCV (12), HPV (13), Ebola (14), Dengue (15), and Zika (15), which contains sequence information on virus proteins with corresponding human protein information and their MiST scores, i.e. their interaction confidence scores (16) and 2) human proteome fasta files containing one protein sequence per gene (16). Mostly, project time was spent covering data processing, support vector machines (SVMs), and deep learning using Python. The group put these concepts in practice through hands-on work with their individual project; the six students chose one of six pathogens (HIV, HCV, HPV, Ebola, Dengue, and Zika) to work on individually. First, students built a dataframe containing their chosen virus-human interactions and split the data frame into a positive dataset with host-pathogen interactions with MiST score >= 0.75 and a negative dataset with proteins of their chosen virus and a randomized human proteins, resulting in 248 positive interactions and 496 negative interactions for Dengue, 89 positive and 140 negative interactions for HCV, 704 positive and 1400 negative interactions for Zika, 93 positive and 186 negative interactions for HIV. The students then applied a global sequence alignment using a Biopython package (17) to check the pattern of the sequence alignments between positive and negative interactions. To predict their virus-human protein interactions, students combined the negative and positive datasets and created a SVM. The students coded separately on personal Jupyter notebooks but shared code through CoLaboratory notebooks and collaborated through project time discussion, screen-sharing, and Slack messaging.

In the first week, the group began by reading three papers on PPI predictive techniques and dissected the various merits of each paper’s methodology before deciding to pursue sequence alignment to demonstrate homology between their protein family and another potentially related member. Next, the students accessed the PPI dataset of host-pathogen PPI data containing virus bait protein, corresponding human prey protein and gene name from PubMed for their chosen pathogen. In their first dataset, they mainly organized the bait and prey sequences and corresponding MiST score in a comprehensive and cohesive format. Each student chose a different virus of six options. Together, they collaboratively processed the primary PPI dataset by isolating their virus’ bait and protein sequence to build their virus-protein dataframe. To close off the week, the instructors introduced the second training dataset consisting of each of the six pathogens’ protein ID and sequences. The students downloaded and utilized fasta files from UniProt that contained the protein ID and sequences for HIV, HCV, HPV Ebola, Dengue, Zika and spent the remainder of project work time understanding the relationship between prey and protein sequences.

To start the second week, students learned about different sequence alignment algorithms. First, the students split each pair of interacting virus bait and human prey depending on the MiST score into positive (MiST >= 0.75) and negative (MiST < 0.75) datasets. After splitting the dataset, the instructors guided the students in constructing a data processing pipeline prior to building their predictive model. To add features of the data, the group utilized a global pairwise alignment algorithm from Biopython (17) to add the sequence alignment scores for each bait and prey pair to the positive and negative dataframes. Guided by their instructors, the group deliberated and decided on features that may serve as potent predictive variables including bait protein length, amino acid counts, and the atomic weight of the bait protein. Finally, the students visualized the distribution of alignment scores for the positive and negative data and evaluated the association.

In the final week, the group began creating and testing their machine learning models. The students combined the positive and negative dataframes and visualized their data through a scatter plot. Next, the group implemented SVMs and collaboratively built their classifier. Students selected an 80% and 20% split for their training and testing data, respectively. Each student first trained their model using their individual virus data. Then, they trained the model using all their virus data to predict the interaction between each SARS-CoV-2 protein and each human protein from the first PPI dataframe they built. The students finetuned the algorithmic parameters, to improve the model’s performance. To visualize the algorithm’s optimal performance, each student built a confusion matrix for the SVM predicting virus-human protein interaction (**Fig 2a-e**) and extracted feature importance in a bar plot (**Fig 2f**). Additionally, students were guided by their instructors to build a convolutional neural network (CNN) for their individual pathogen.

**Fig 2.**
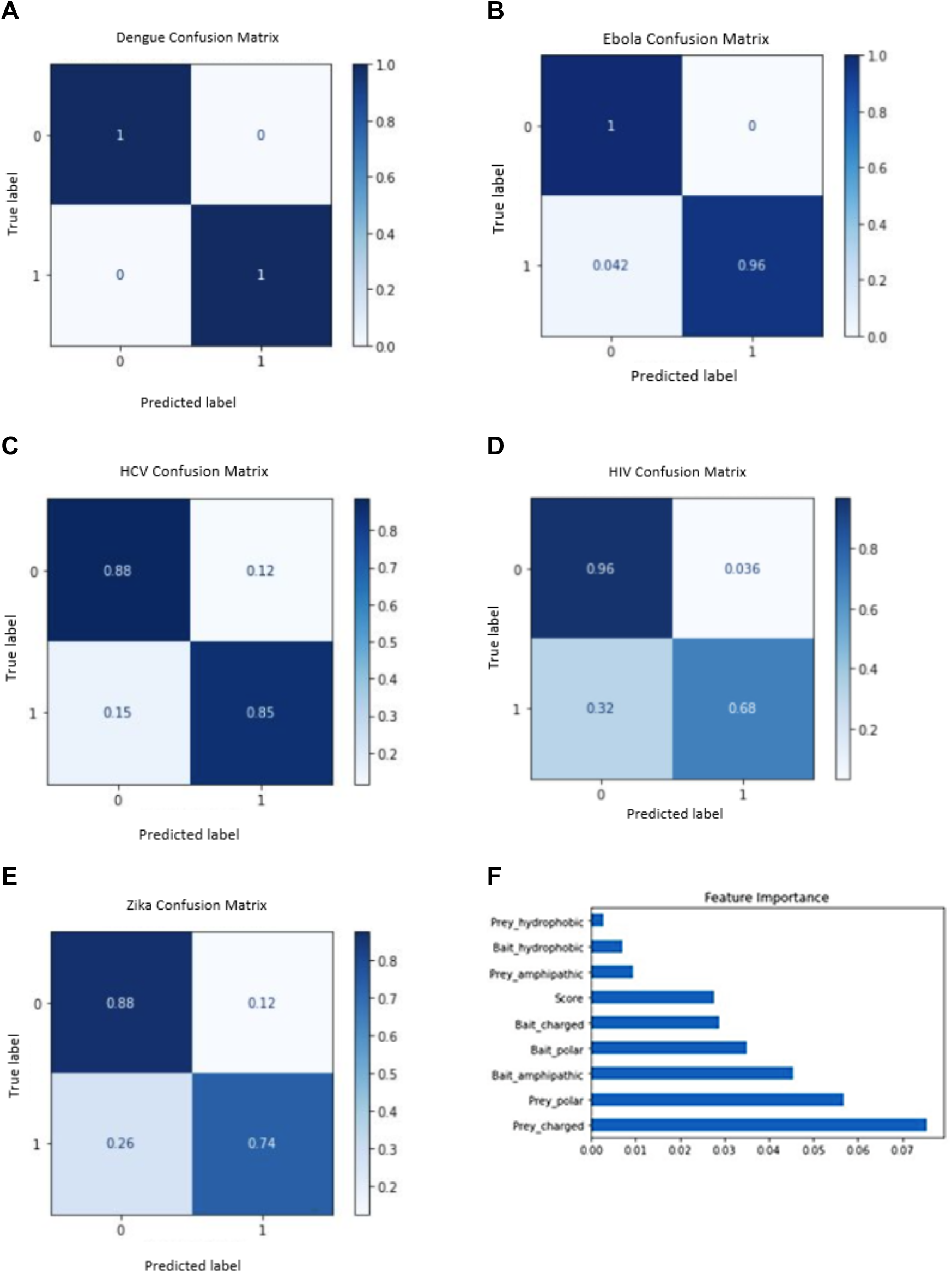
AI and Proteomics: COVID-19 Protein-Protein Interactions (PPI) Predictions. A-E: Confusion matrices visualizing the individual SVM models’ performances. Evaluated the true and predicted virus-human protein interactions for Dengue, Ebola, HCV, HIV, and Zika viruses respectively. **F: Bar plot of feature importance extracted from Ebola virus trained SVM on SARS-CoV-2 virus data.** The prey proteins with a net positive or negative charge had the highest feature importance.

Finally, the students spent the remainder of the last week investigating the implications of their research. The instructors presented ways that this type of data can be leveraged for drug discovery and repurposing whereupon the students discussed the implications of their experimental results on the COVID-19 pandemic.

Code availability, data, slides, and figures can be found <u>here</u>.

#### Project 3: AI for Imaging Data - Computer Vision for COVID-19 Chest X-Ray Classification

Students learned to implement machine learning models that can classify COVID-19 cases in chest x-ray images. The dataset was curated from publicly available chest x-ray images datasets (18,19) and contained 438 images from patients diagnosed with COVID-19 as well as 438 images from patients without pathologic findings, labeled as ‘no finding’ (**Fig 3a-b**). The instructors introduced concepts in data processing, computer vision, and deep learning using Python. The students applied these concepts in a hands-on project where they first visualized and evaluated the dataset then trained a convolutional neural network (CNN) to identify COVID-19 cases. The students coded separately on personal CoLab notebooks but collaborated and debugged together to the extent possible through online video discussions and screen-sharing.

**Fig 3.**
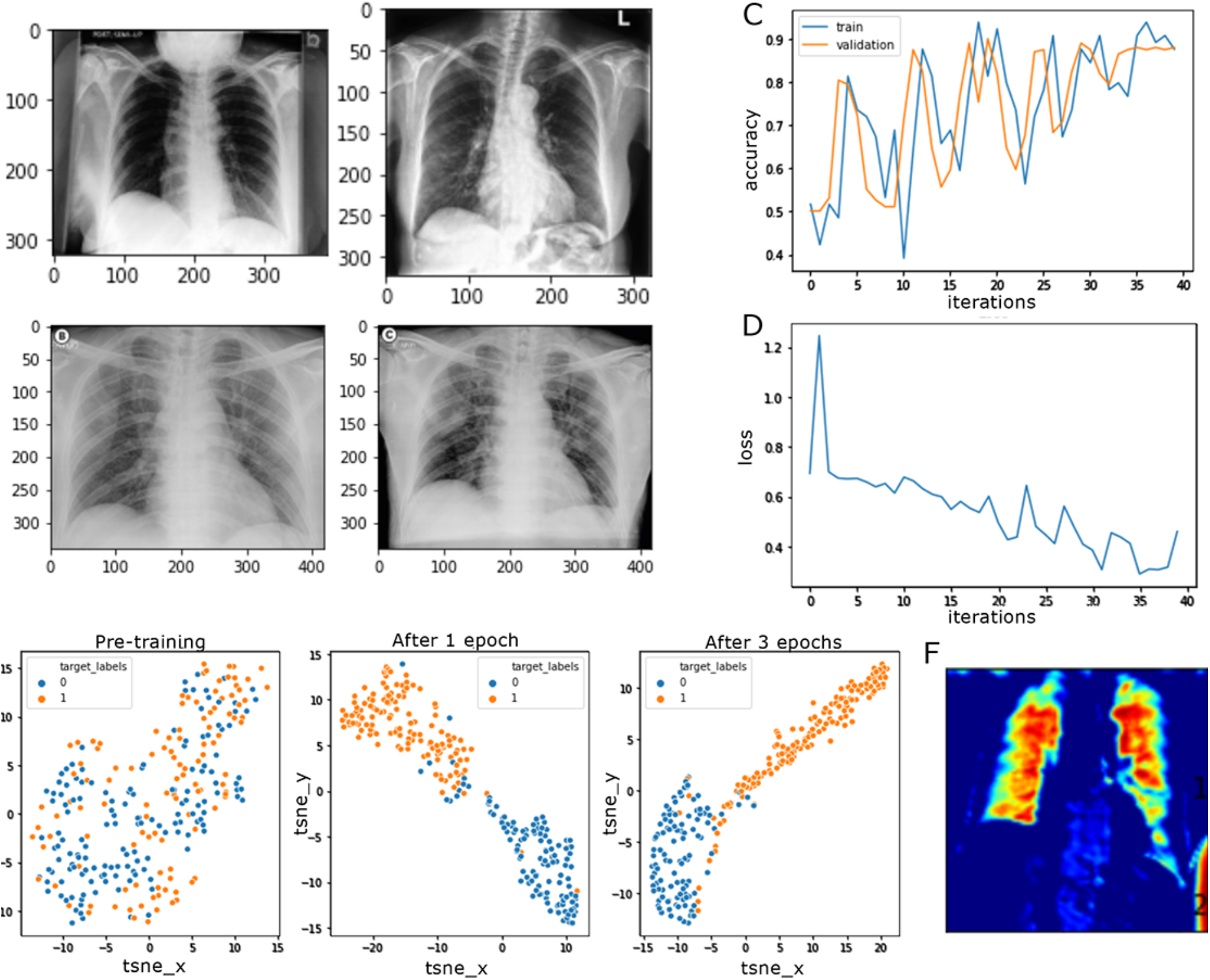
AI for Imaging Data. **A. Subset of no finding images.** Courtesy of Pranav Rajpurkar and Andrew Ng (CheXpert)(19). **B. Subset of COVID-19 images.** Credit: Lim et al (J Korean Med Sci, 2020)(20) and Dr Domenico Nicoletti (Radiopaedia.org, rID: 74724)(21). **C. Accuracy graph during model training. D. Loss graph during model training. E. TSNE visualization of model training over time.** Blue represents no finding images while orange represents COVID-19 images. **F. Grad-CAM interpretation of CNN features.** Red in the heatmap represents greater importance while blue represents the least.

The group began by examining the dataset through exploratory data analysis. They experimented with different data visualization approaches and packages to plot the distribution of attributes in the dataset. Using the visualizations, the students discovered potential biases in the dataset such as skewed gender distribution and different x-ray views for the COVID-19 images versus the ‘no finding’ images. The students also learned to use dataframe manipulations to survey subsets of the datasets. Together, they identified image features, such as lung region opacity and body outlines, that could be useful or potentially problematic for the classification task.

Next, the students developed fully-connected and convolutional neural networks (CNN) using PyTorch to perform binary classification. The instructors provided a starting code framework that guided the students in constructing a typical data processing and machine learning pipeline. The students completed the code by implementing missing core sections. They tracked model training speed as well as accuracy and loss curves to gain insight into the model training process (**Fig 3c-d**). The students also calculated metrics, such as F1 scores, and visualized latent space features to evaluate the model (**Fig 3e**). By discussing these analyses, the students identified areas where the model is performing poorly (e.g. incorrectly classifying COVID-19 images) and formulated hypotheses for potential reasons, such as the large variability in the COVID-19 images.

In the final week, the students branched out to work on follow-up ideas. For this half of the project, the students were largely self-driven to explore ideas or questions that interested them. The instructors helped the students work through code issues and brainstormed solutions. The follow-up ideas included iteratively improving CNN models by tuning hyperparameters and training other types of classifiers (SVM, regression) using features extracted by a pre-trained DenseNet model. One particular challenge the students tackled was model robustness. Using gradient class activation maps to provide interpretation of model decisions, they had observed that the CNN models were detecting regions outside of the lung (**Fig 3f**). This became a focus for model improvement. During breaks, the instructors also briefly introduced interesting papers and recent results in AI vision research. The group discussed challenges of interpretable decision-making and model brittleness, which complemented the tasks they were working on.

#### Project 4: Latent Variable Modeling of COVID-19 Metagenome Transcriptomic Profiles

Students applied dimensionality reduction techniques to investigate natural stratification of SARS-CoV2+ patient subgroups based on host transcriptomic response and viral coinfection status. Previous analyses of metagenomic sequencing data from upper airway samples of 238 patients revealed a diminished innate immune response in patients positive for SARS-CoV2 through differential expression analysis, gene set enrichment analysis, and *in silico* estimation of cell type proportions (22). We hypothesized decomposition of metagenomic next generation RNA sequencing (mNGS) to reveal separable clusters of patient subgroups. In our approach, we applied both supervised and unsupervised decomposition methods to analyze structural patterns inherent to the data.

First, the students accessed this publicly available data and performed exploratory analysis on the study cohort of 94 patients who tested positive for SARS-CoV2 by gold standard clinical PCR. Mick et. al quantified the abundance of 15,900 host genes and 275 viruses by RNA sequencing and reference based alignment. Students applied principal component analysis (PCA) to each dataset and qualitatively inspected emergence of clusters based on covariates such as SARS-CoV2 viral load, gender, and age. In this unsupervised analysis the students experienced how sample outliers can skew variance and cause inflation of PCA components. In order to account for possible non-linear structure within the data, the students also trained and tested autoencoders on both the viral coinfection dataset and the gene expression dataset.

Evaluation of viral coinfection embeddings led the students to hypothesize a correlation between binary coinfection status and SARS-CoV2 viral load. The students reasoned that patients with additional viral infection(s) may be generally more susceptible to SARS-CoV2 replication due to alteration of immune response dynamics under coinfection conditions. Patients were stratified into 2 groups: those with measurable alternative viral load, and those with sole viral metagenome alignment to SARS-CoV2. Welch’s t-test was applied to the average reads-per-million (rPM) of SARS-CoV2 between the two groups, and a significant difference below an error tolerance of 0.05 was found (p=1.947*10-4) (**Fig 4a**).

**Fig 4.**
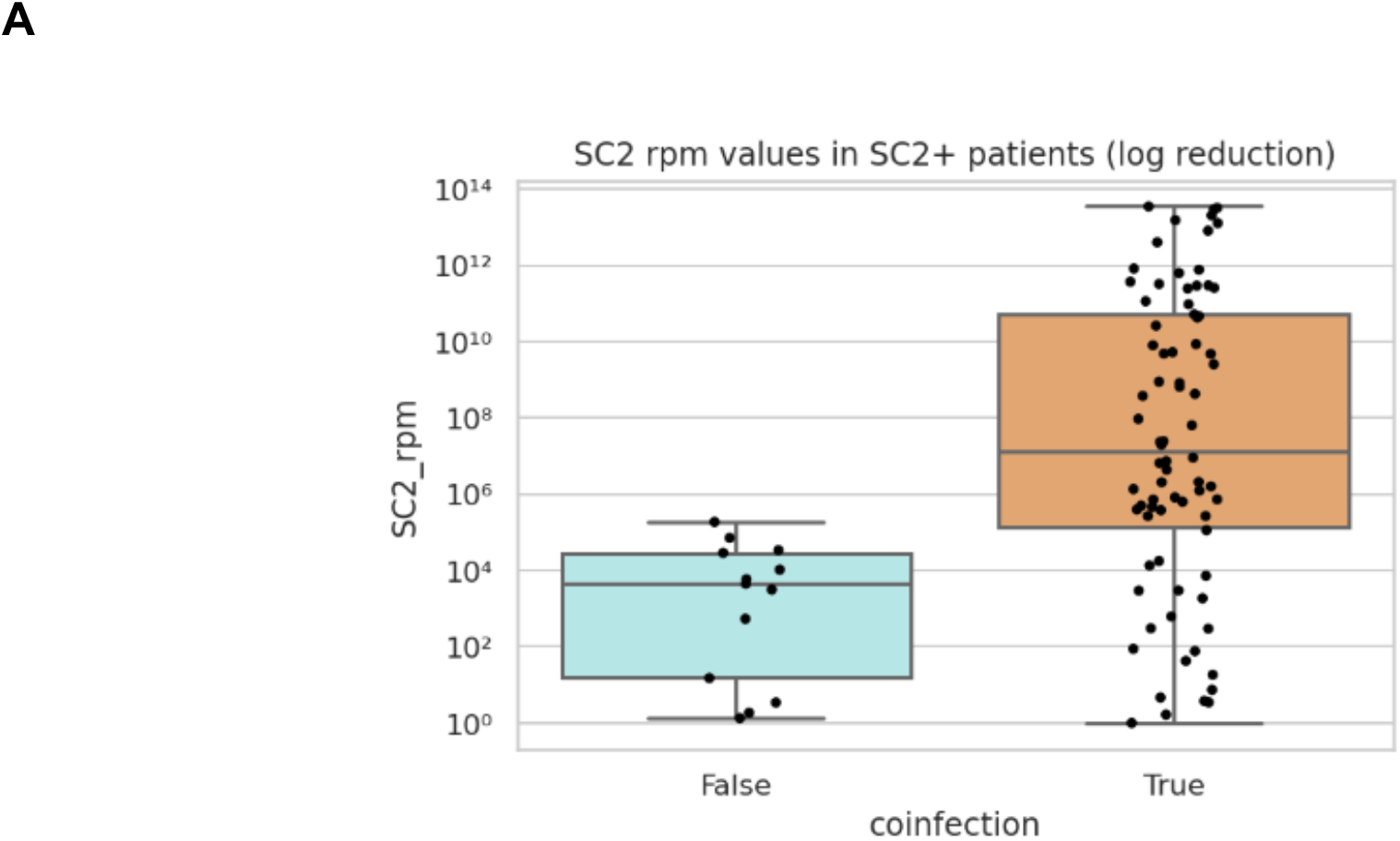
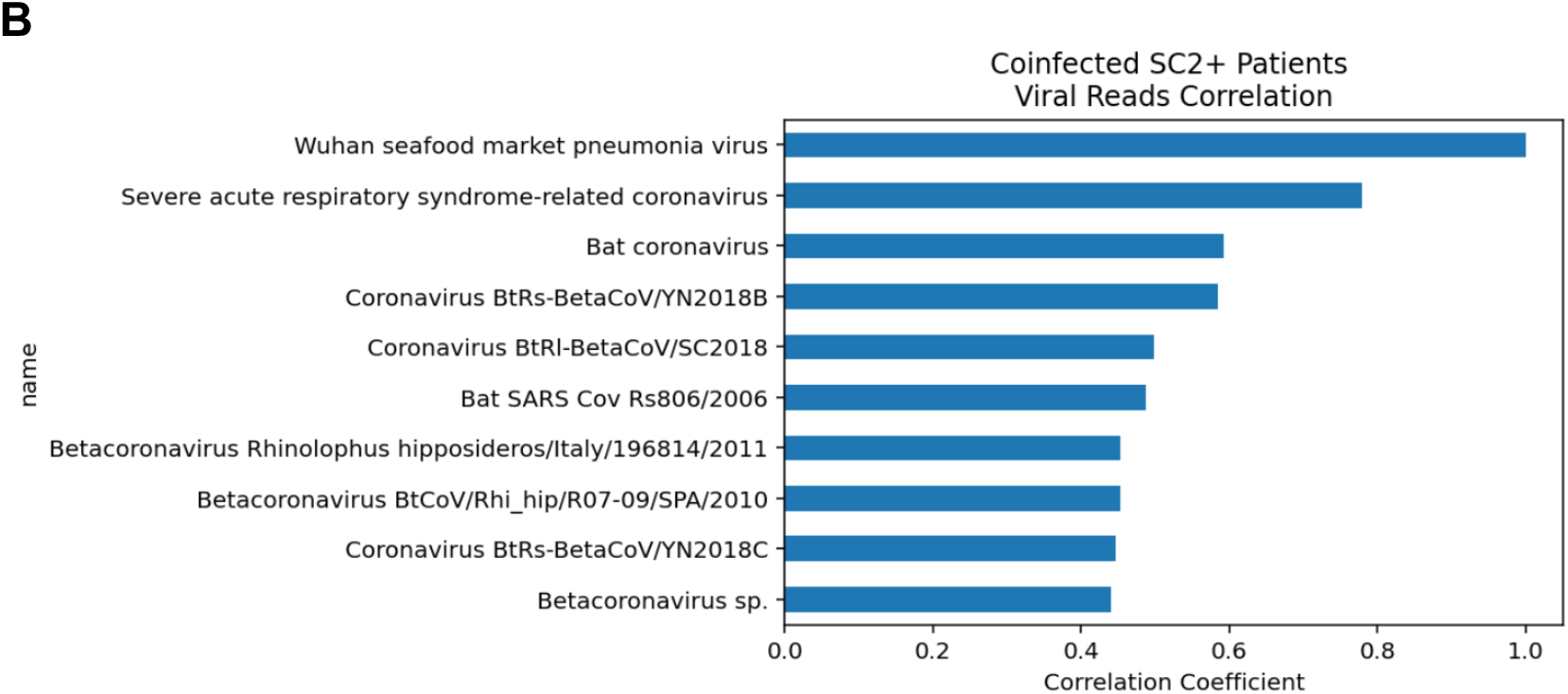
Latent Variable Modeling of COVID-19 Metagenome Transcriptomic Profiles. **A. Boxplot of SARS-CoV2 rPM by coinfection status.** Reads-per-million of SARS-CoV2 stratified by patients with alternative (orange) and no (blue) detectable viruses. **B. Correlation coefficients between reads of SARS-CoV2 and coinfected viruses.** Highly correlated viruses are coronaviruses.

The students followed up this result with an analysis for confounding variables. In group discussion, we posited that samples with a large SARS-CoV2 viral load cause a higher frequency of alignment errors to evolutionarily similar genomes. To test this, the students inspected correlation coefficients between SARS-CoV2 and alternative, coinfection viral read abundance. The most correlated viruses were all coronaviruses, lending evidence to the hypothesis that higher SARS-CoV2 viral load results in a higher proportion of reference alignment errors to genomically similar viruses (**Fig 4b**). This experience underscored the adage, “correlation does not imply causation,” lending a valuable lesson to the analysis of SARS-CoV2+ patient subgroups.

#### Project 5: PredictCOVID - AI and Electronic Medical Record (EMR) data

Using real world data from (https://www.kaggle.com/einsteindata4u/covid19), students learned how to apply AI to (A) predict whether a patient is COVID positive or negative and (B) predict the severity of the COVID infection (i.e. admission into the general ward, semi-intensive care unit, or intensive care unit). The dataset included 5,644 patients as well as COVID-19 test results, patient age quantile, hospital admission ward, and various laboratory results from blood tests, urine tests, and pathogen tests. Students were introduced to the benefits and drawbacks of publicly available data, such as sources of bias and the need for intensive data preprocessing before the data can be utilized. Students took different approaches to data cleaning and imputation of missing values and evaluated the performance of machine learning models on varied input data, including evaluating metrics of accuracy, area under the curve (AUC), and distribution of false negatives and positives (**Fig 5a**).

**Fig 5.**
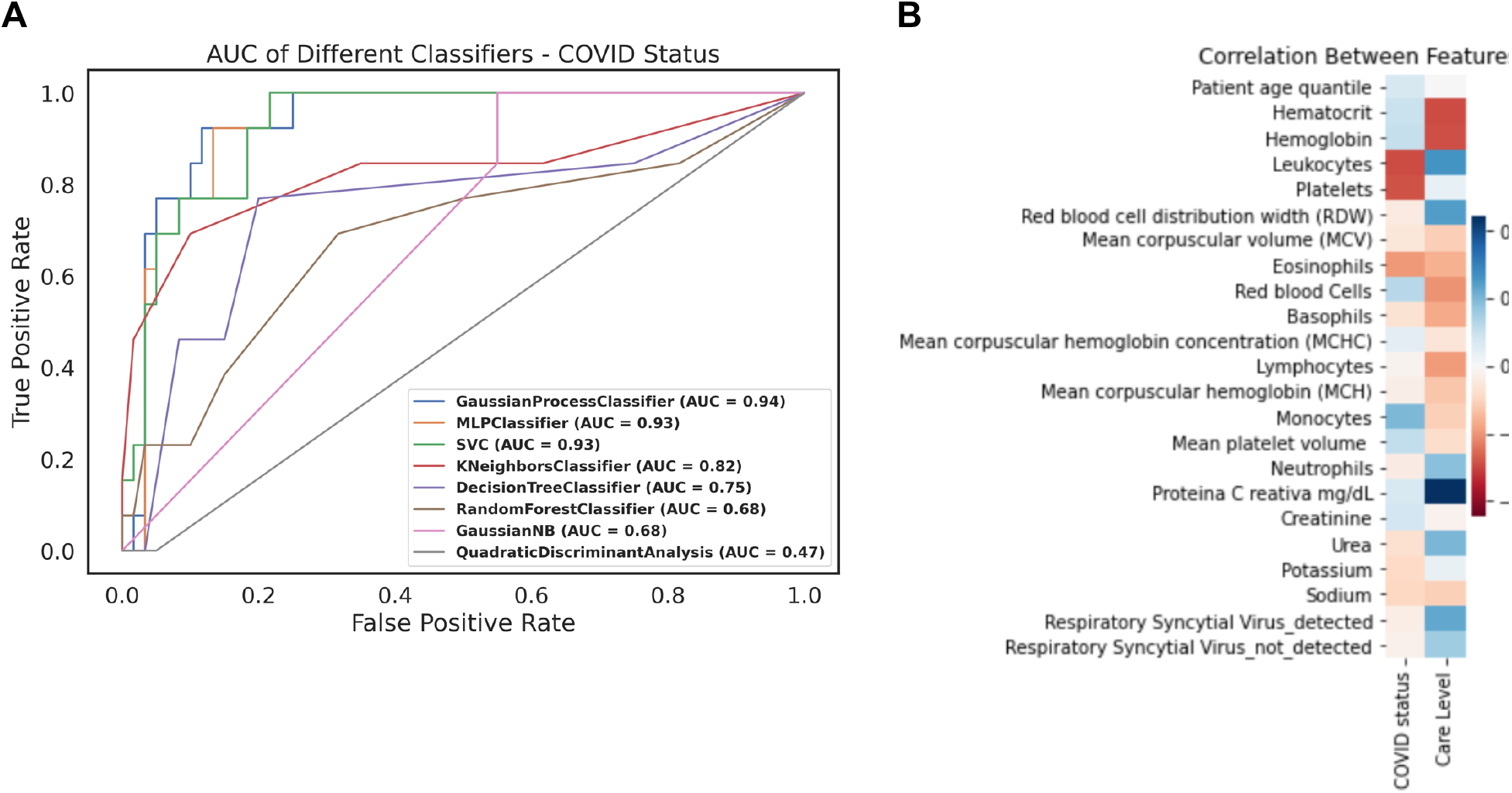
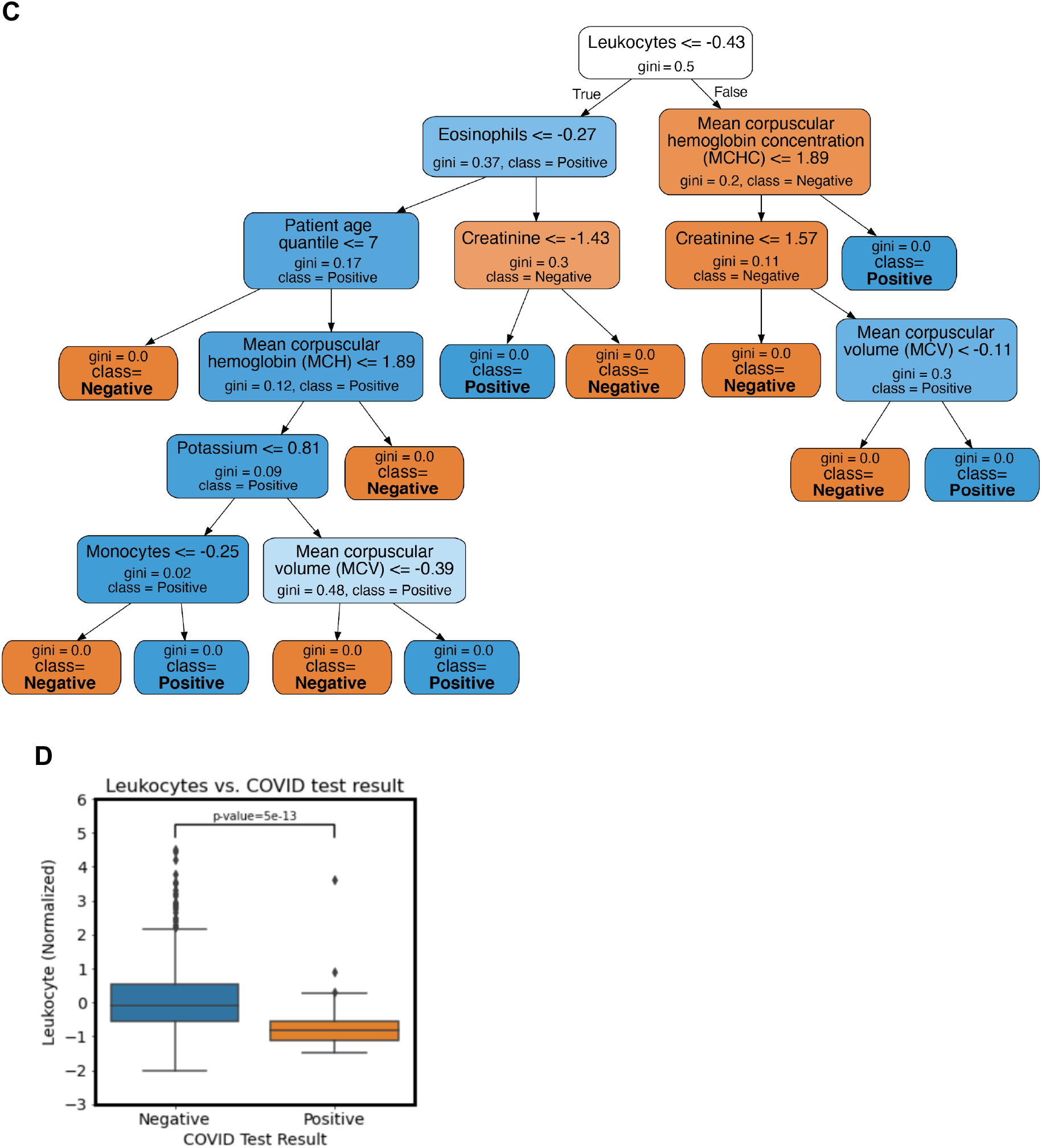
PredictCOVID: AI for deriving insights from healthcare data. A. Area Under the Curves (AUCs) of different machine learning classifiers for predicting COVID status from patient data. AUC was one of the metrics used by the students to evaluate the performance of their machine learning models.**B. Decision tree classifier for COVID status.** Features used in decision making in each level of the tree are shown, with COVID status (positive/negative) at the leaves of the tree. **C. Heatmap of the correlation between clinical features and COVID status or Care Level.** Blue represents positive correlation while red represents negative correlation. **D. Boxplot of normalized leukocyte laboratory values and COVID status.** Leukocyte laboratory values negatively correlated with COVID test results.

Additionally, students were asked to go above and beyond to apply their findings to translational applications. For example, students were asked to critically evaluate the cost of false negatives (spreading COVID-19, not receiving treatment on time, worse outcomes) and false positives (waste of limited resources) in respect to patients and outcomes, and applying this evaluation to the decision of a model. Students were also asked to perform covariate analyses to determine feature importance and apply back to their understanding of clinical relevance and application (**Fig 5b,c**). One finding that the group reported was that leukocytes were heavily negatively correlated with COVID test results (**Fig 5d**). Lastly, the group summarized their findings and recommendations for future plans to the entire group as well as the limitations and biases in the data (i.e. single location, limited follow-up, missing data).

### Pre- and Post- Surveys

Analysis of the survey data revealed significant shifts in some of the students’ responses from the Pre- to Post- survey. Specifically, at the end of the program, there were significantly more students who reported that they know how to clean data before using it in machine learning algorithms (MWU test, adjusted p-value <0.001), and know how to evaluate and apply machine learning algorithms (MWU test, adjusted p-values <0.001) (**Fig 6**). More students also reported knowing people in AI who are people of color (MWU test, unadjusted p-value = 0.014, adjusted p-value = 0.285) and women (MWU test, unadjusted p-value = 0.044, adjusted p-value = 0.877), feeling confident in questioning the media about AI (MWU test, unadjusted p-value = 0.015, adjusted p-value = 0.297), and knowing about careers that use AI (MWU test, unadjusted p-value = 0.037, adjusted p-value = 0.743); however, these increases were only of nominal significance. Survey responses of students in the 2020 virtual program and those of students in the 2019 commuter program for questions asked in both years were found not to differ significantly (Mann Whitney U test, unadjusted and adjusted p-values > 0.05) (**Supplementary Table 1**). Students of the 2020 virtual were also no less likely to recommend the AI4ALL program to peers than the students who attended the 2019 in-person program (Mann Whitney U test, unadjusted p-value = 0.044, adjusted p-value = 0.872) (**Supplementary Figure 1**).

**Fig 6.**
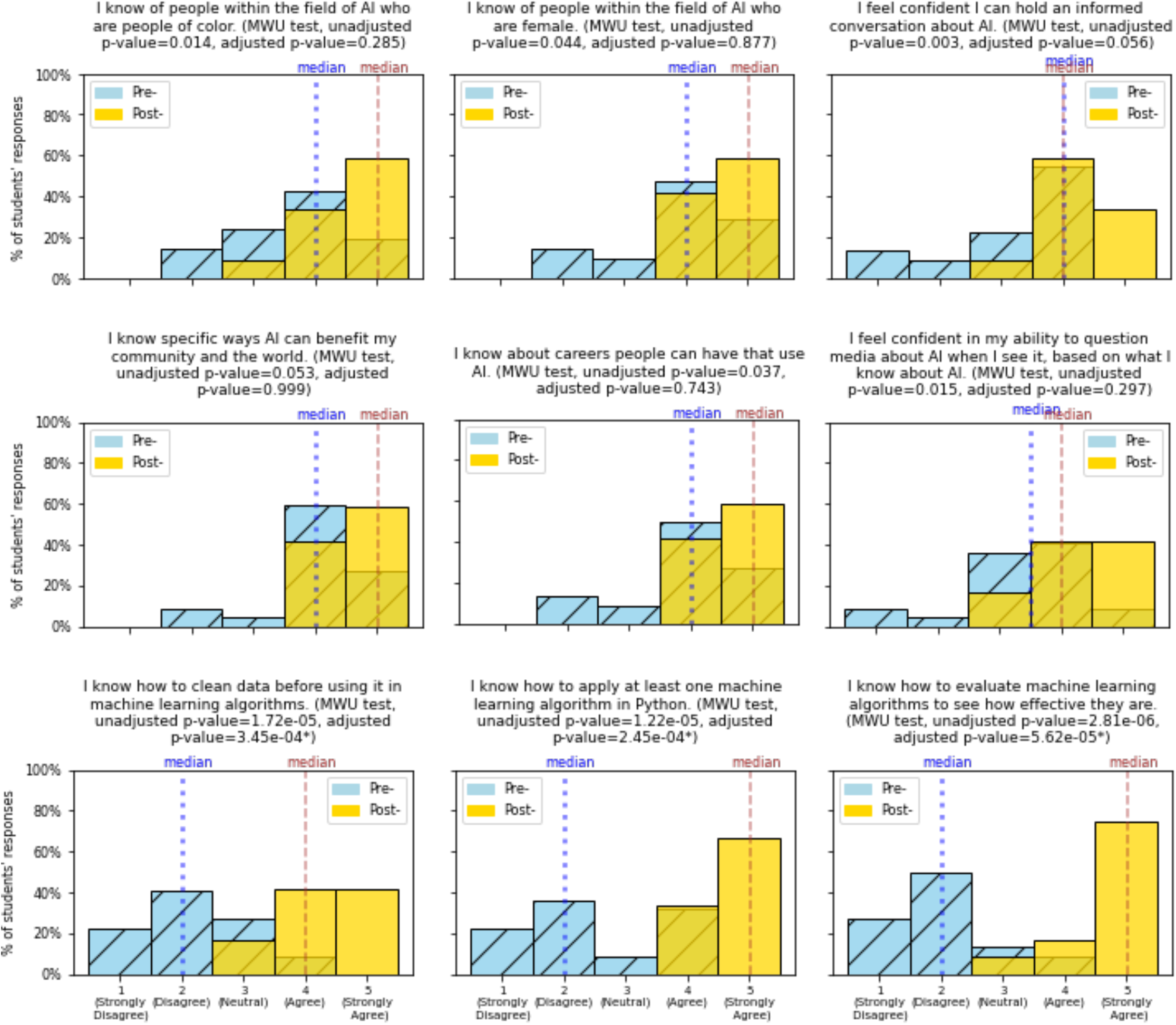
Pre-post survey results. Histograms of students’ responses to pre-program and post-program survey questions, with median values. Mann Whitney U tests were performed to compare pre- and post-survey responses, with adjusted p-values below significance threshold of 0.05 given an asterisk (*).

Moreover, the graduate students, post-doctoral scholar and alumni TA’s who worked with our students on their research projects shared their observations that the students:

○ Grew in their abilities to think critically, form hypotheses, and design executable experiments
○ Learned how to develop a variety of different ML models
○ Fortified their collaborative skills and technical proficiency in Python
○ Found exploratory data analysis to be valuable as it challenged the participants to view scientific inquiry in an open-ended way that deviated from traditional classroom experiences
○ Learned how to analyze the results and how they relate to the original research questions
○ Experienced the immediate relevancy of AI approaches to current problems in the COVID-19 pandemic
○ Demonstrated their understanding and interpretation of not only AI but also the application of AI to medicine, public health, and clinical decision making

## Discussion

Diverse representation is needed not just in the data for Artificial Intelligence (AI), but also in the people working and leading in the field of AI. Since 2019, UCSF AI4ALL has engaged students from backgrounds historically underrepresented in AI in order to promote greater diversity and inclusion in this field. In 2020, through a variety of interactive virtual real-time sessions and experiences held during a three-week period, our program allowed students to interact with a diverse set of role models in AI and learn about how AI can be used to advance health.

Furthermore, students gained experience in coding, working with data, and AI by participating in one of our meaningful hands-on research projects that applied AI to understanding, classifying, or predicting COVID-19.

Students’ survey responses demonstrated their feeling significantly more familiar with working with data and evaluating and applying machine learning algorithms at the end of our 2020 virtual program. There was also a nominally significant increase in the students’ knowing people in AI who are from historically underrepresented groups, their confidence in discussing AI, and their awareness of careers in AI. While the format of the 2020 program differed from 2019, with the 2020 program taking place online instead of in-person due to the pandemic, students’ survey responses from both years were comparable.

Despite the success of our virtual training program, there were some limitations to having a program take place entirely online, including the lack of in person interactions and the need for reliable internet connection. Nevertheless, the ability to engage young students in AI and the opportunity to contribute to diverse representation in this field make holding our program in any format worthwhile.

We have learned that it is possible to deliver virtually an AI curriculum to young high school students that provides them with an engaging and impactful experience. Through our virtual program, we were able to connect with students from around the country and involve teaching assistants and faculty from outside the Bay Area and from other institutions. We were also able to give students who are located far from AI training programs a chance to become involved bringing the goal of increasing diversity in AI a little closer to reality.

## Author Contributions

TO and MS designed and co-directed the program, performed analysis of program survey data, outlined and wrote the manuscript. JW, IK, and JB led and described Project 1. ZN and IK led and described Project 2. IC, TG, and JY led and described Project 3. WC, RB, and CS led and described Project 4. AT and BV led and described Project 5. JB, TC, WC, SE, TG, KL, AT, and DMT developed curriculum materials and led instructional sessions. All authors discussed results, provided critical feedback and contributed to the final manuscript.

## AI4ALL Student Cohort 2020

Arhana Aatresh^18^, Githika Annapureddy^18^, Ami Baid^18^, Qixin Dai^18^, Esha Gohil^18^, Yomn Hammad^18^, Ishika Hazra^18^, Vienna Huang^18^, Valerie Kwek^18^, Marissa Lee^18^, Angela Liu^18^, Neha Naveen^18^, Ava Paikeday^18^, Sonica Prakash^18^, Srihita Ramini^18^, Yamuna Rao^18^, Neeharika Ravi^18^, Mihika Rayan^18^, Brenda Samano^18^, Jessica Sanchez^18^, Anooshkha Shetty^18^, Arely Sun^18^, Riya Tadi^18^, Monica Trinh^18^, Gianna Yan^18^, Joyce Yang^18^, Caroline Yoon^18^, Fiona Zhang^18^, and Hannah Zhuang^18^.

## Acknowledgements

The authors would like to acknowledge everyone in the AI4ALL organization, including Tess Posner, Olga Russakovsky, Fei Fei Li, Valerie Allen, Beth McBride, Jonathan Garcia, and Thalea Torres, and our colleagues in the Bakar Computational Health Sciences Institute, including Angela Jackson and Sharat Israni, for their encouragement and support. We would also like to acknowledge Aeshah Al-Nagdawi, Gaia Andreoletti, Matthew Hancock, Andrew Jan, Chakrapani Kalyanaraman, Idit Kosti, Brian Le, Hunter Mills, Boris Oskotsky, Ben Rubin, Wren Saylor, Gundolf Schenk, Victoria Turner, Rohit Vashisht, and Katharine Yu for their work to support our Program; Sourav Bandyopadhyay, Sergio Baranzini, Atul Butte, Gabriela Fragiadakis, Julian Hong, Michael Keiser, Sara Murray, Matthew Spitzer, and Ida Sim for participating as our Faculty Guest Speakers; Marilyn Ritchie for her Keynote presentation; Maya Gonzalez, Nishant Jain, Anooshree Sengupta, and Carolyn Wang for participating in our Undergraduate Students Panel; Stella Belonwu, Emmalyn Chen, and Caroline Warly-Solsberg for participating in our Graduate Students Panel; Sarah Aerni, Tiffany Chen, Sam Gross, Katie Planey, and Alex Morgan for participating in our Bio in AI Industry Panel; Arohee Bhoja, Tia Jain, Christine Li, and Sachi Parikh for participating in our UCSF AI4ALL Alumni Panel; Bill Lindstaedt for leading our students in a Personal Growth Session; the Altschuler and Wu Lab, the Giacomini Lab, the Keiser Lab, the Kroetz Lab, the Krogan Lab, and the Witte Lab at UCSF for their support of our program; Pranav Rajpurkar and Andrew Ng for granting us permission to include images from CheXpert; and Edna Rodas and Amber Nolan for Administrative Support.

## References

1. 8 Helpful Everyday Examples of Artificial Intelligence [Internet]. IoT For All. 2020 [cited 2021 Feb 2]. Available from: https://www.iotforall.com/8-helpful-everyday-examples-of-artificial-intelligence

2. Yu K-H, Beam AL, Kohane IS. Artificial intelligence in healthcare. Nat Biomed Eng. 2018 Oct;2(10):719–31.

3. Fröhlich H, Balling R, Beerenwinkel N, Kohlbacher O, Kumar S, Lengauer T, et al. From hype to reality: data science enabling personalized medicine. BMC Med. 2018 Aug 27;16(1):150.

4. Noor P. Can we trust AI not to further embed racial bias and prejudice? BMJ. 2020 Feb 12;368:m363.

5. Adamson AS, Smith A. Machine Learning and Health Care Disparities in Dermatology. JAMA Dermatol. 2018 Nov 1;154(11):1247.

6. West SM, Whittaker M, Crawford K. Discriminating Systems: Gender, Race, and Power in AI [Internet]. AI Now Institute; 2019 Apr [cited 2021 Jan 5] p. 33. Available from: m https://ainowinstitute.org/discriminatingsystems.html

7. Dong E, Du H, Gardner L. An interactive web-based dashboard to track COVID-19 in real time. Lancet Infect Dis. 2020 May 1;20(5):533–4.

8. Akaike H. Fitting autoregressive models for prediction. Ann Inst Stat Math. 1969;21:243–7.

9. Chatfield C. The Holt-Winters Forecasting Procedure. J R Stat Soc Ser C Appl Stat. 1978;27(3):264–79.

10. Kröse B, Krose B, Smagt P van der, Smagt P. An introduction to Neural Networks. 1993.

11. Jäger S, Cimermancic P, Gulbahce N, Johnson JR, McGovern KE, Clarke SC, et al. Global landscape of HIV-human protein complexes. Nature. 2011 Dec 21;481(7381):365–70.

12. Ramage HR, Kumar GR, Verschueren E, Johnson JR, Von Dollen J, Johnson T, et al. A Combined Proteomics/Genomics Approach Links Hepatitis C Virus Infection with Nonsense-Mediated mRNA Decay. Mol Cell. 2015 Jan 22;57(2):329–40.

13. Eckhardt M, Zhang W, Gross AM, Von Dollen J, Johnson JR, Franks-Skiba KE, et al. Multiple Routes to Oncogenesis Are Promoted by the Human Papillomavirus-Host Protein Network. Cancer Discov. 2018 Nov;8(11):1474–89.

14. Batra J, Hultquist JF, Liu D, Shtanko O, Von Dollen J, Satkamp L, et al. Protein Interaction Mapping Identifies RBBP6 as a Negative Regulator of Ebola Virus Replication. Cell. 2018 Dec 13;175(7):1917-1930.e13.

15. Shah PS, Link N, Jang GM, Sharp PP, Zhu T, Swaney DL, et al. Comparative Flavivirus-Host Protein Interaction Mapping Reveals Mechanisms of Dengue and Zika Virus Pathogenesis. Cell. 2018 Dec 13;175(7):1931-1945.e18.

16. Breuza L, Poux S, Estreicher A, Famiglietti ML, Magrane M, Tognolli M, et al. The UniProtKB guide to the human proteome. Database J Biol Databases Curation [Internet]. 2016 Feb 19 [cited 2021 Feb 16];2016. Available from: https://www.ncbi.nlm.nih.gov/pmc/articles/PMC4761109/

17. Cock PJA, Antao T, Chang JT, Chapman BA, Cox CJ, Dalke A, et al. Biopython: freely available Python tools for computational molecular biology and bioinformatics. Bioinformatics. 2009 Jun 1;25(11):1422–3.

18. Cohen JP, Morrison P, Dao L. COVID-19 Image Data Collection. ArXiv200311597 Cs Eess Q-Bio [Internet]. 2020 Mar 25 [cited 2021 Feb 3]; Available from: http://arxiv.org/abs/2003.11597

19. Irvin J, Rajpurkar P, Ko M, Yu Y, Ciurea-Ilcus S, Chute C, et al. CheXpert: A Large Chest Radiograph Dataset with Uncertainty Labels and Expert Comparison. Proc AAAI Conf Artif Intell. 2019 Jul 17;33(01):590–7.

20. Lim J, Jeon S, Shin H-Y, Kim MJ, Seong YM, Lee WJ, et al. Case of the Index Patient Who Caused Tertiary Transmission of Coronavirus Disease 2019 in Korea: the Application of Lopinavir/Ritonavir for the Treatment of COVID-19 Pneumonia Monitored by Quantitative RT-PCR. J Korean Med Sci [Internet]. 2020 Jan 8 [cited 2021 Feb 25];35(6). Available from: https://doi.org/10.3346/jkms.2020.35.e79

21. COVID-19 pneumonia | Radiology Case | Radiopaedia.org [Internet]. [cited 2021 Feb 25]. Available from: https://radiopaedia.org/cases/covid-19-pneumonia-7

22. Mick E, Kamm J, Pisco AO, Ratnasiri K, Babik JM, Calfee CS, et al. Upper airway gene expression differentiates COVID-19 from other acute respiratory illnesses and reveals suppression of innate immune responses by SARS-CoV-2. medRxiv. 2020 May 22;2020.05.18.20105171.

