## Supplementary Materials for "Nurturing diversity and inclusion in AI in Biomedicine through a virtual summer program for high school students"

### Supplementary Tables and Figures

| Survey Question | 2020 Pre-Median | 2019 Pre-Median | MWU test unadjusted p-value | MWU test adjusted p-value | 2020 Post-Median | 2019 Post-Median | MWU test unadjusted p-value | MWU test adjusted p-value |
|---|---|---|---|---|---|---|---|---|
| I know of people within the field of AI who are people of color. | 4 | 3 | 0.332 | 0.999 | 5 | 5 | 0.532 | 0.999 |
| I know of people within the field of AI who are female. | 4 | 3.5 | 0.617 | 0.999 | 5 | 5 | 0.755 | 0.999 |
| I know about careers people can have that use AI. | 4 | 4 | 0.242 | 0.999 | 5 | 5 | 0.969 | 0.999 |
| I feel confident I can hold an informed conversation about AI. | 4 | 3 | 0.370 | 0.999 | 4 | 4 | 0.777 | 0.999 |
| I know specific ways AI can benefit my community and the world. | 4 | 5 | 0.068 | 0.136 | 5 | 5 | 0.307 | 0.999 |

**Supplementary Table 1. Comparison of 2020 and 2019 survey responses.** Medians and Mann Whitney U (MWU) test p-values for comparison of survey responses of students in the 2020 virtual program and those of students in the 2019 commuter program for questions asked both years.

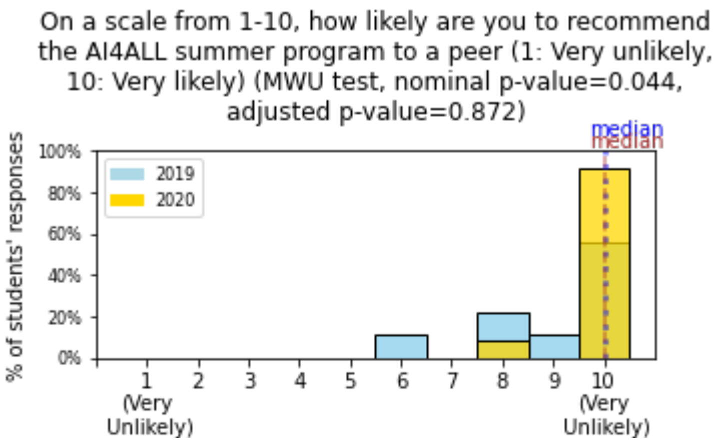


**Supplementary Figure 1. Likeliness to recommend program in 2019 vs 2020.** Histogram of 2019 and 2020 students' responses to how likely they were to recommend the AI4ALL summer program to a peer. Mann Whitney U tests were performed to compare responses.
